# Intravenous anti-Aβ immunotherapy acutely increases cerebral amyloid angiopathy and vascular damages in APOE4 mice

**DOI:** 10.64898/2026.02.09.704876

**Authors:** Philip Pikus, Gracie S. Healey, Elizabeth Xia, Giorgi Shautidze, Neha Siddapureddy, Yichien Lee, Christopher Albanese, Olga Rodriguez, R. Scott Turner, G. William Rebeck

## Abstract

Anti-Aβ immunotherapies for Alzheimer’s Disease (AD) have high rates of amyloid-related imaging abnormalities (ARIA), an adverse side effect with markedly higher rates in *APOE4* carriers. We developed a mouse model of ARIA centered on human *APOE3* and *APOE4* genotypes with amyloidosis (5xFAD transgene) and microglia tagged with green fluorescent protein (from the CX3CR1 promoter). We measured acute changes following a single intravenous treatment with 3D6 anti-Aβ immunotherapy. Across 82 mice, *APOE4* mice showed stepwise reductions in the number of plaques from one to ten days, with significant reductions in the subiculum (48%) and thalamus (40%) at ten days. There was no significant reduction in *APOE3* mice. There was a concomitant significant increase in deposition of cerebral amyloid angiopathy (CAA) in *APOE4* mice at one (76%) and three (51%) days in leptomeningeal vessels. The increased CAA correlated with a significant 189% increase in Aβ within microglia of *APOE4* (but not *APOE3*) mice at one day. Smooth muscle actin staining showed significant 58% reduction near CAA. MRI analysis revealed a significant 32% increase in microhemorrhages ten days following treatment. These data demonstrate an *APOE4*-specific redistribution of parenchymal amyloid to CAA by 3D6 within days, leading to increased vascular damages associated with ARIA.

## Introduction

Until 2021, no FDA-approved disease modifying therapies for Alzheimer’s Disease (AD) existed. This represented a major challenge, as an estimated 50 million people suffer from dementia worldwide, with AD contributing to 60-70% of these cases ^1^. The prevalence of AD is expected to grow further in the coming years as the population ages. Several novel immunotherapies targeting distinct forms and epitopes of amyloid-beta (Aβ) have been shown to clear Aβ deposits both in preclinical models ^2^ and in large-scale clinical trials, with these trials showing that these treatments can slow cognitive decline over an approximately 18-month period in patients with mild cognitive impairment or mild dementia due to AD ^3,4^.

The excitement surrounding these novel therapies has been offset by high rates of an adverse side effect known as amyloid related imaging abnormalities (ARIA). ARIA occurs in up to a third of patients receiving these therapies ^5^, with subgroups of ARIA-E (edema) and ARIA-H (microhemorrhage) associated with the leakage of fluid or blood products into the brain parenchyma, respectively. While many patients are asymptomatic with lesions only appearing on MRI, others experience headaches, nausea, mental status changes, gait disturbances, seizures, and in rare cases, coma or death ^5^. The risk of ARIA is highest immediately following the first infusion of antibody, with close to 70% of ARIA occurring following the first two infusions across clinical trials ^6,7,8^. The underlying mechanism of ARIA remains unknown, with risk factors including APOE4 genotype, cerebral amyloid angiopathy (CAA), dose of treatment, and antithrombotic use ^5^. ARIA is correlated with plaque clearance in meta-analyses across multiple antibodies ^6^. There is an urgent need to better understand the mechanisms that contribute to ARIA, particularly in the acute stages of treatment.

Aβ plaques precede the clinical symptoms of AD and contribute to neuroinflammation, cell death, cerebral vascular damage, and dementia, forming the basis of the amyloid cascade hypothesis ^9^. Aβ can deposit in the brain in two primary forms: parenchymal plaques which are composed primarily of Aβ_42_, and amyloid that deposits along the walls of the cerebral vasculature composed primarily of Aβ_40_ ^10^. CAA can occur independently or as a co-pathology in about 30% of AD cases ^11,12^. In CAA, Aβ deposits around the smooth muscle layer of arteries and arterioles, often in the leptomeningeal regions of the brain ^10^. In addition to being a co-pathology with AD, CAA is associated with white matter damage, intracerebral hemorrhage, and related inflammation (CAA-ri) ^5,6^. Neuropathological studies have found high levels of CAA in several individuals who received active or passive anti-amyloid immunotherapies ^13–15^, leading to the hypothesis that an increase in CAA may occur with these therapies ^16^.

*APOE* genotype is the strongest risk factor for late onset AD. Existing in three different alleles, each *APOE* ε4 allele increases risk of AD, while each *APOE* ε2 allele is neuroprotective ^17^. The *APOE4* genotype increases Aβ deposits (both parenchymal and CAA), while the *APOE2* genotype results in less plaque deposition but paradoxically increased CAA ^18,19^. *APOE4* genotype is also the strongest risk factor for ARIA, with rates increasing between 2- to 5-fold for both ARIA-E and ARIA-H across clinical trials ^5,20^. This association both presents a risk for patients receiving immunotherapies and precludes many *APOE4* individuals from initiating treatment altogether. Given that existing microhemorrhages seen on baseline MRI (often reflective of advanced CAA) increase risk of ARIA and are a contraindication for these therapies ^21,22^, many of these patients are either unable to pursue treatment or cautioned against it by their providers. There are no established mechanisms for why ARIA occurs at such increased rates in *APOE4* individuals ^23^. Given that *APOE4* homozygotes make up 10-20% of AD patients despite making up only 2-3% of the total population ^24,25^, it is critical to better understand the role of APOE genotype in ARIA to more safely treat these patients.

Numerous preclinical studies have established that anti-Aβ immunotherapies clear plaques (see reviews) ^2,26–28^, and chronic treatment can clear vascular Aβ deposits ^29–31^. However, few studies have measured the acute effects of these therapies on levels of CAA, and studies investigating levels of CAA following more intermediate treatment schedules of passive immunotherapy have produced mixed results ^30,32^. No studies have addressed what happens to parenchymal or vascular Aβ deposits in the immediate period following initiation of IV immunotherapy, the period when ARIA occurs most frequently. Our study addresses this question by measuring levels of plaques and CAA in the brain parenchyma and leptomeninges at different acute timepoints following IV treatment with 3D6 anti-Aβ immunotherapy. We use a model incorporating human *APOE3* or *APOE4*, investigating both the cellular changes associated with anti-Aβ immunotherapies and the incidence of ARIA-H by MRI. We find that a single dose of 3D6 immunotherapy causes *APOE4-*specific increases in plaque clearance, CAA deposition, and smooth muscle cell damage in only one to ten days. By ten days, *APOE4* mice show increased brain microhemorrhages, consistent with the increased rates of ARIA-H in *APOE4* individuals.

## Results

### Development of novel mouse model of ARIA

While many mouse models exist to study amyloidosis and AD, few models combine Aβ deposition, measures of neuroinflammation, and APOE genotype. We generated a novel mouse model incorporating the 5xFAD model of amyloidosis ^33^, green-fluorescent microglia expressed on the Cx3Cr1 promoter ^34^, and knock-in human *APOE3* or *APOE4* genotypes (Figure 1A) ^35^. We injected 3D6 intravenously (IV) into the lateral tail vein to measure neuropathological changes at 1-,3-, and 10-days post-treatment (Figure 1B). These mice develop amyloid pathology after two months of age, with significant plaque deposition by four months of age (Supplemental Figure 1), at which point they exhibit aggressive amyloidosis in the subiculum, entorhinal cortex, and regions of the thalamus (Figure 1C). CAA is observed sporadically throughout the brain tissue under these conditions (Figure 1D). To validate that 3D6 reached the brains and reacted with plaques, we used a fluorescently conjugated form of 3D6 (3D6-647); by three days, this tagged 3D6 was present throughout the brain and co-localized prominently with Aβ deposits (Figure 1E).

**Figure 1.**
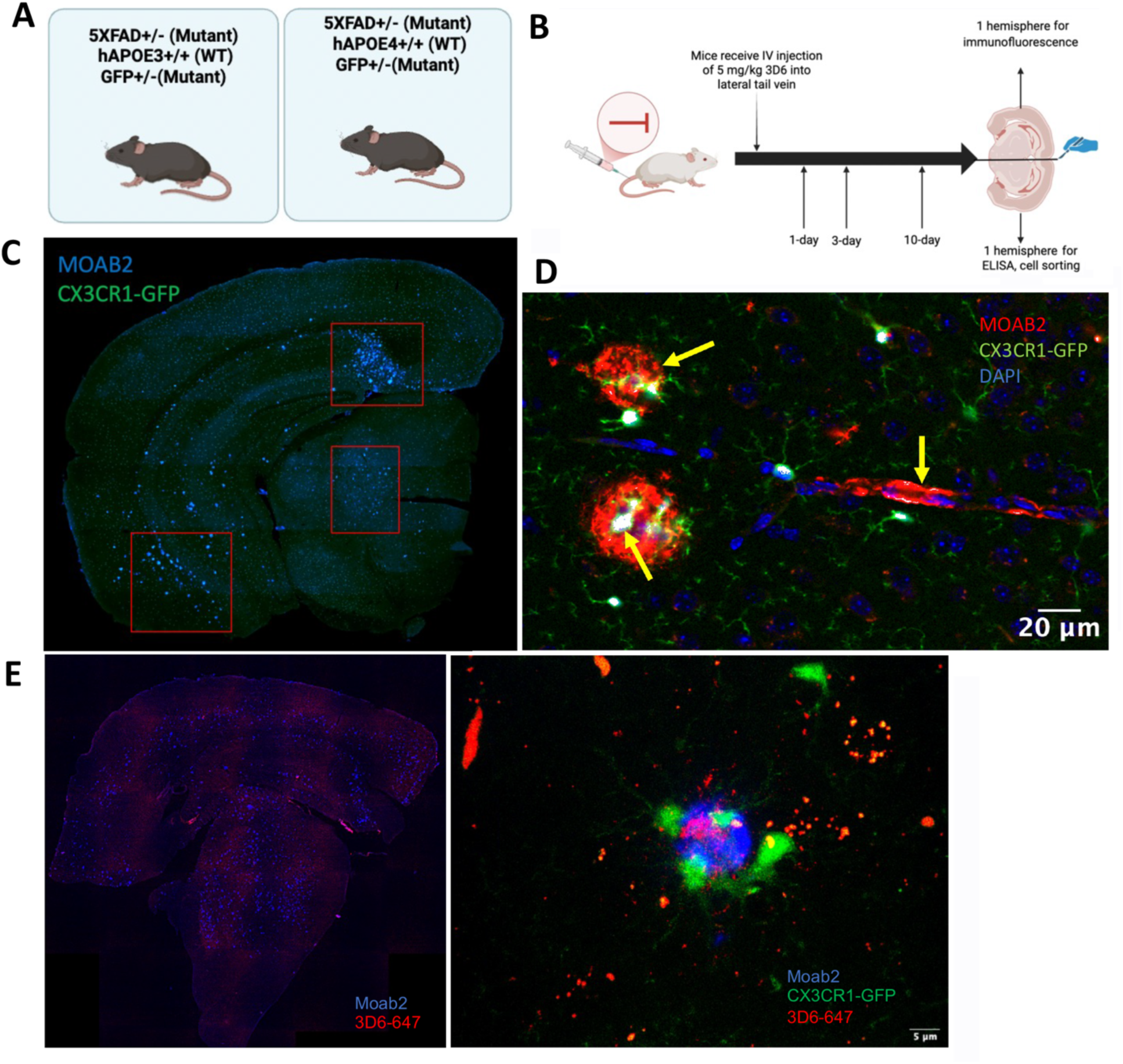
Developing a novel mouse model of IV anti-Aβ immunotherapy in *APOE3* and *APOE4* transgenic mice. A. Mice of desired genotype: 5xFAD^+/−^, CX3CR1^GFP/-^, *APOE3*^+/+^ or 5xFAD^+/−^, CX3CR1^GFP/-^, *APOE4*^+/+^. B. Treatment paradigm and timeline with 4-month-old female and male mice receiving IV injections of 3D6 or IgG1 isotype control antibody and euthanized for experiments at 24-hours, 72-hours, and 10-days post-injection. C. Brain hemisphere showing distribution of Aβ plaques in the subiculum, entorhinal cortex, and thalamus (boxed) in 4-month-old untreated mouse. D. Higher magnification image showing Moab2 staining of Aβ both at amyloid plaques and CAA, and GFP-fluorescent microglia in the subiculum in a 3D6 treated mouse 72-hours post-injection. E. Fluorescently conjugated 3D6 antibody (3D6-647) reached the brain (left) and reacted with plaques (right).

### Plaque clearance occurs preferentially in APOE4 mice

We assessed amyloid clearance in plaque-rich regions by counting the number of plaques and percent area coverage after immunostaining with the Moab2 anti-Aβ antibody. A representative image of the subiculum from an IgG1 isotype control mouse with marked amyloidosis is shown (Figure 2A). As expected, *APOE4* mice had a higher baseline plaque-load compared with *APOE3* mice at 4-months-of-age. This trend was consistent across brain regions, with significantly more plaques in the *APOE4* subiculum (110%, p<0.01) (Figure 2B). Mice were administered 3D6 intravenously at this age, and we measured the number of plaques and the percent brain area coverage of amyloid at 24-hours, 72-hours, and 10-days post-injection. In *APOE4* mice 10 days post-treatment, the number of plaques were significantly decreased (Figure 2C) in the subiculum (48%, p<0.01) and thalamus (40%, p<0.05), with lesser (non-significant) reductions at 24 and 72 hours (Figure 2D). Clearance of plaques was not observed in any region at these time points in *APOE3* mice (Figure 2G), indicating an acute 3D6-mediated clearance of plaques that occurs preferentially in *APOE4* mice. There was a similar but less significant effect in region-specific amyloid load. In *APOE4* mice, there was a significant decrease in percent area coverage between mice in the 72-hour and 10-day timepoints in the entorhinal cortex (p<0.05). In all three brain regions, there were non-significant decreases comparing 10 days of treatment to the IgG controls (subiculum: 62%, p=0.08; thalamus: 53%, p=0.12; entorhinal cortex: 50%, p=0.35) (Figure 2F).

**Figure 2.**
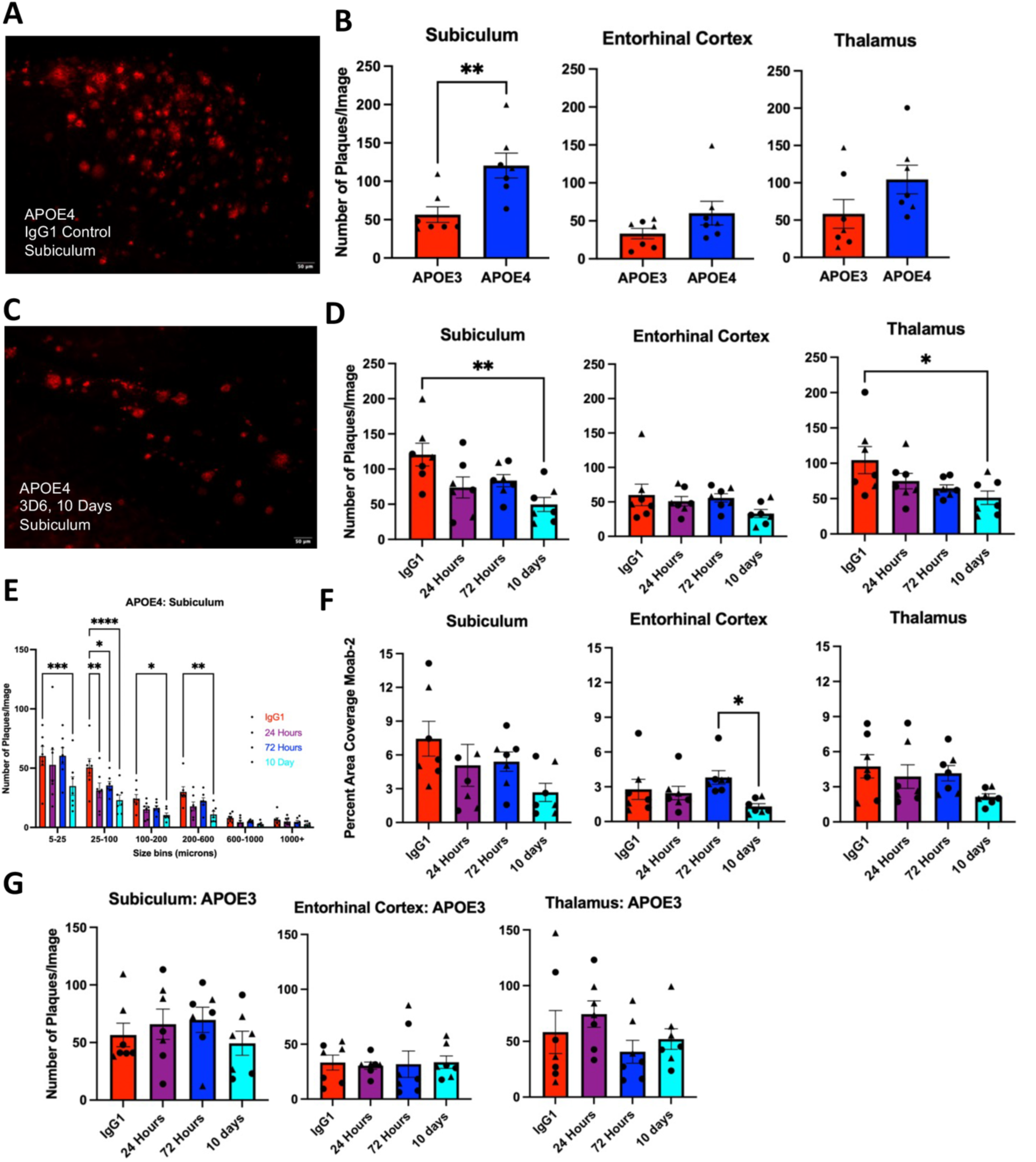
IV 3D6 immunotherapy acutely clears a portion of plaques by 10-days post-treatment in a region-specific manner, preferentially in APOE4 mice. A. Representative image of plaque distribution in the subiculum from an IgG1 isotype control treated mouse. B. Student’s t-tests measuring mean ± SEM number of plaques/image in regions of interest in *APOE3* vs. *APOE4* mice at baseline (IgG isotype control) (N=7 mice/group, *p<0.05, **p<0.01). C. Representative image of plaque distribution in the subiculum from a 3D6-treated mouse (10-days) showing clearance of a portion of plaques. D. One-way ANOVAs with multiple comparisons measuring mean ± SEM number of plaques/image in *APOE4* mice at different timepoints in different regions following IV 3D6 immunotherapy (N=7 mice/group, *p<0.05, **p<0.01). E. Two-way ANOVA with multiple comparisons measuring mean ± SEM number of plaques/image in different size bins of Aβ plaques in the subiculum. Bins are 5-25 μm^2^, 25-100 μm^2^, 100-200 μm^2^, 200-600 mm^2^, 600-1000 μm^2^, and over 1000 μm^2^ (N=7 mice/group, *p<0.05, **p<0.01, ***p<0.001, ****p<0.0001). F. One-way ANOVA with multiple comparisons measuring mean ± SEM percent area coverage of Moab-2 (Aβ) at different timepoints in regions of interest in *APOE4* mice (N=7 mice/group, *p<0.05). G. One-way ANOVA with multiple comparisons measuring mean ± SEM number of plaques/image in *APOE3* mice at different timepoints in the subiculum, entorhinal cortex, and thalamus showing lack of changes (N=7 mice/group).

### 3D6 clears amyloid deposits of smaller and medium sizes

Given that 3D6 acutely cleared plaques in *APOE4* mice, we measured whether there were differences in clearance between plaques of different sizes. We divided amyloid deposits into different size bins: 5-25 μm^2^ (small “sub-plaque” deposits), 26-100 μm^2^ (small plaques), 101-200 μm^2^ (small-medium plaques), 201-600 μm^2^ (medium plaques), 601-1000 μm^2^ (large plaques), and >1000 μm^2^ (very large deposits). In the subiculum, there was a decrease in the number of both small and medium-sized plaques by day 10 (Figure 2E). There was no statistically significant reduction in the number of plaques of larger sizes, although there were fewer plaques at baseline in these size bins. In the entorhinal cortex, reduction of plaque numbers was observed in the 25-100 μm^2^ range at 10-days post-treatment, and in the thalamus, there was only a significant decrease in smaller deposits, similar to, but less significant than, the changes in the subiculum (Supplemental Figure 2).

### 3D6 increases CAA in both the parenchyma and leptomeninges in APOE4 mice

There are low levels of CAA present at baseline (IgG1-treatment control). In 3D6-treated mice, the CAA is easily apparent around alpha-smooth muscle cells at leptomeningeal vessels, (Figure 3A) and in the parenchyma (Figure 3B). We tested for changes in the number of loci of CAA in the leptomeninges at different timepoints following IV 3D6 immunotherapy. Representative images are shown for leptomeningeal CAA in an *APOE3* mouse (left) and an *APOE4* mouse (right) treated with 3D6 for 24-hours (Figure 3C). There were no *APOE* genotype effects on the number of CAA vessels at baseline (Figure 3D). There was a significant increase in the number of vessels impacted by CAA at 24-hours (76%, p<0.001) and 72-hours (51%, p<0.05) in the leptomeninges in *APOE4* mice, with levels returned close to baseline by day 10 (Figure 3D). These changes were not seen in *APOE3* mice (Figure 3D). Similarly, in the parenchyma, there were increased numbers of CAA vessels at 72-hours (75%, p<0.01) in 3D6-treated *APOE4* mice, with no corresponding changes in *APOE3* mice (Figure 3E). These data show an *APOE4-*specific increase in CAA levels within one and three days after anti-amyloid antibody treatment.

**Figure 3.**
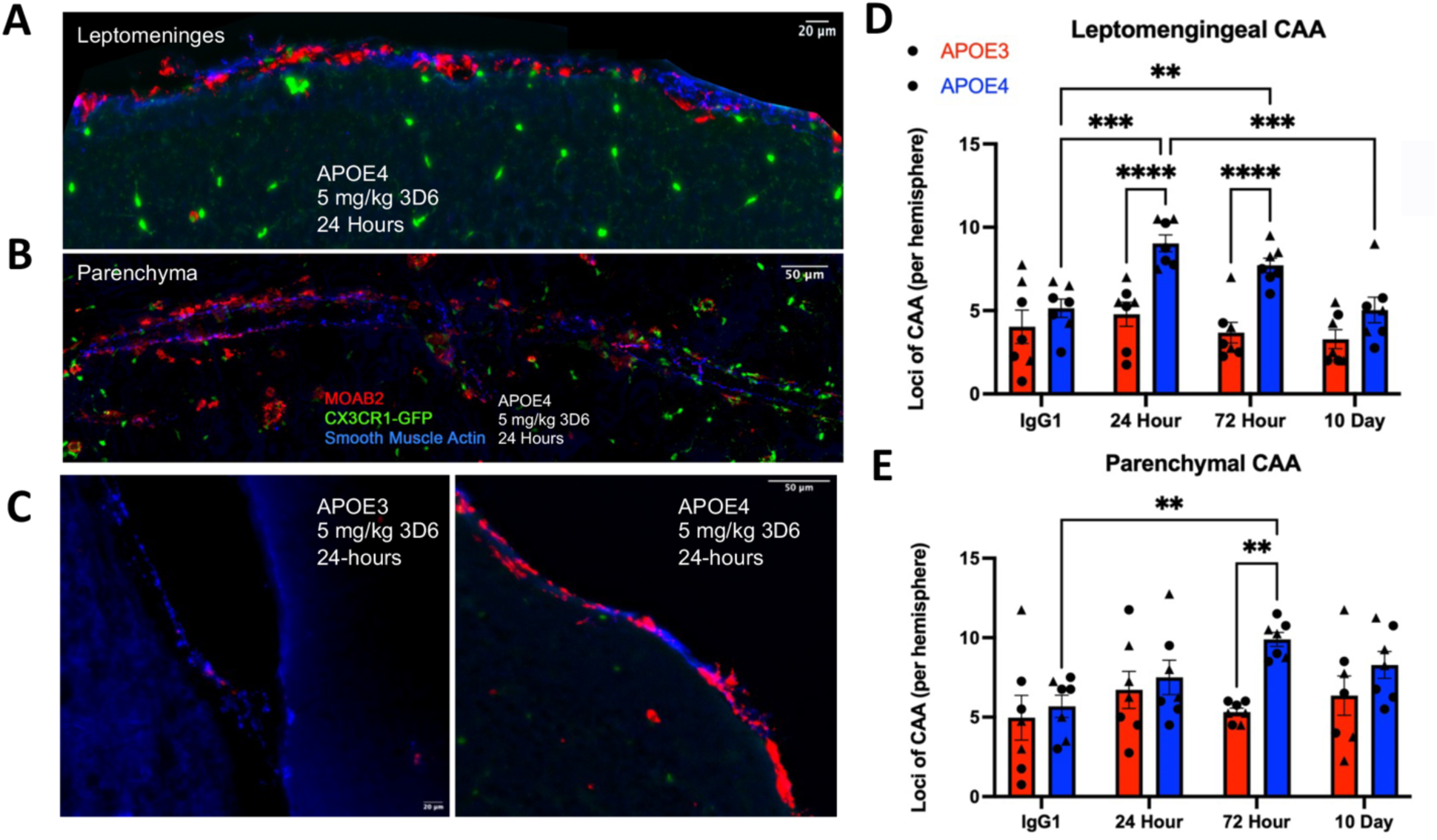
IV 3D6 immunotherapy acutely and transiently increases levels of CAA in both leptomeningeal and parenchymal vessels in *APOE4* but not *APOE3* treated mice. A. CAA accumulation around a leptomeningeal vessel stained with alpha-smooth muscle actin in a 3D6-treated *APOE4* mouse (24-hours). B. CAA accumulation around a parenchymal vessel in a 3D6-treated *APOE4* mouse (24-hours). C. Leptomeningeal vessel stained with alpha-smooth muscle actin with little CAA in an *APOE3* treated mouse at 24-hours (left) vs leptomeningeal vessel containing significant CAA in an *APOE4* 3D6-treated mouse at 24-hours (right). D. Quantification of leptomeningeal CAA in *APOE3* vs. *APOE4* mice using two-way ANOVA with multiple comparisons (N=7 mice/group, *p<0.05, **p<0.01, ***p<0.001, ****p<0.0001). CAA loci are defined by the number of vessels per brain hemisphere affected by CAA. E. Quantification of parenchymal CAA in *APOE3* vs. *APOE4* mice using two-way ANOVA with multiple comparisons (N=7 mice/group, *p<0.05, **p<0.01, ***p<0.001, ****p<0.0001).

### 3D6 increases the number and length of CAA deposits in APOE4 mice

Based on the increased incidence of CAA in *APOE4* mice, we tested different features of this CAA: the number of individual deposits and the length of the leptomeningeal CAA. These measures enabled us to determine whether new small deposits were forming, or whether existing CAA was lengthening. CAA can exist either as small (Figure 4A) or long CAA deposits (Figure 4B). There was no difference in the total number of CAA deposits or the length of CAA at baseline between *APOE3* and *APOE4* mice. However, there was a significant increase in the number of CAA deposits in *APOE4* mice at 24-hours (147%, p<.0001) and 72-hours (86%, p<0.01), mirroring the number of vessels affected (Figure 3). There was no effect of 3D6 on CAA deposits in *APOE3* mice (Figure 4C). Next, we measured the size of these CAA deposits that were forming in *APOE4* mice. The number of CAA deposits increased in all smaller/medium size bins at 24-hours post-injection. At 72-hours there was only a significant increase in deposits in smaller size bins, and at 10-days there was only a significant increase in the number of deposits at the smallest size bin (5-10 μm^2^) (Figure 4D). There was again no significant difference in the total length of CAA deposits per vessel between *APOE3* and *APOE4* mice at baseline. With treatment, there was only a significant increase in length at 72-hours between *APOE3* and *APOE4* mice (51%, p<0.001), with no change in length at 24 hours or 10 days (Figure 4E). Together these data indicate a substantial increase in the number of deposits of CAA that occurs acutely following treatment in *APOE4* mice, with only modest lengthening of CAA at 72 hours.

**Figure 4.**
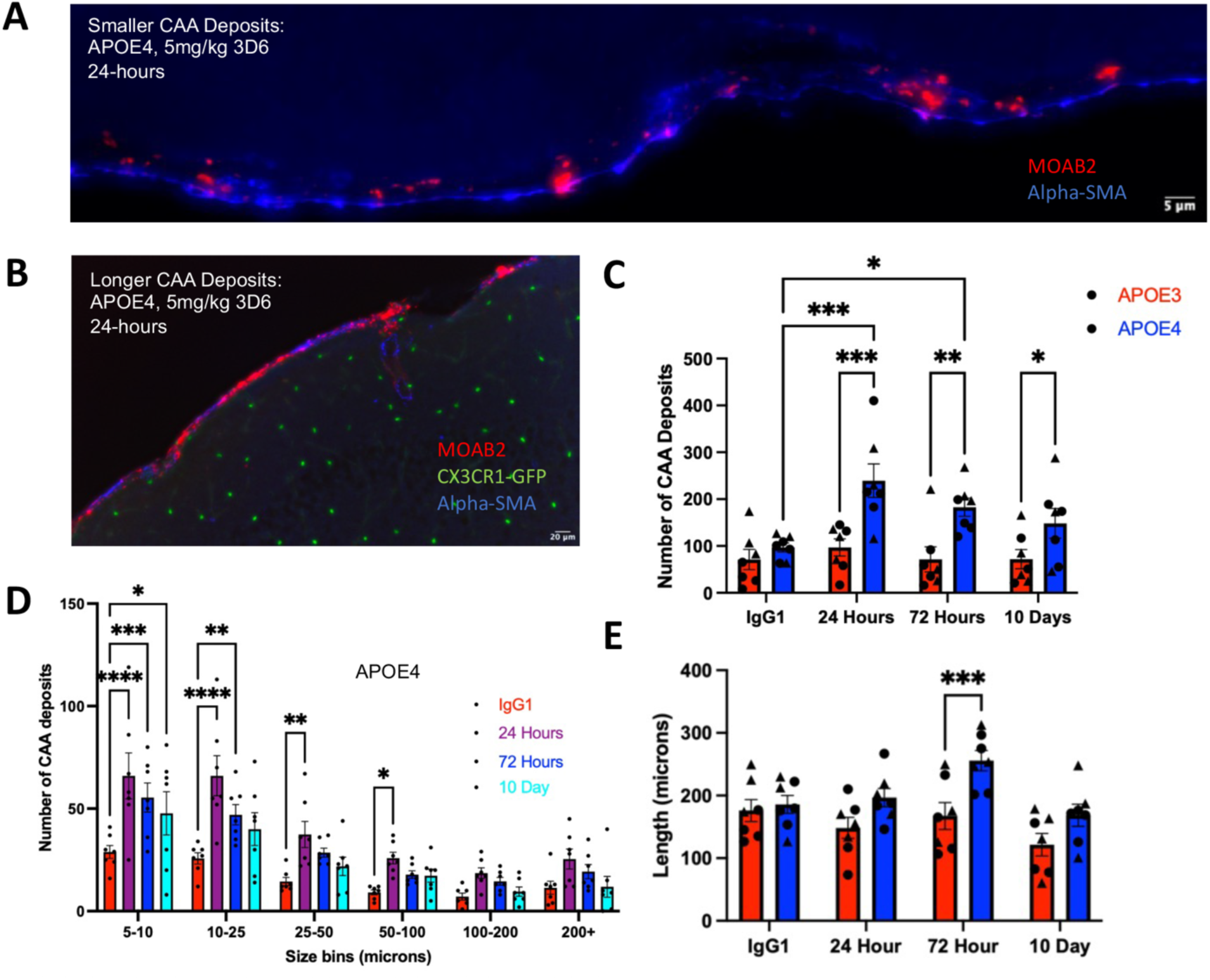
3D6 increases number of CAA deposits while only modestly increasing the length of CAA. A. Representative image showing smaller deposits of CAA along a leptomeningeal vessel stained with alpha-smooth muscle actin. B. Representative image of longer CAA deposition along leptomeningeal vessel. C. 2-way ANOVA with multiple comparisons measuring mean ± SEM number of total CAA deposits in *APOE3* vs. *APOE4* mice at different points. D. 2-way ANOVA with multiple comparisons measuring number of CAA deposits ± SEM in different size bins (5-10, 10-25, 25-50, 50-100, 100-200, 200+ μm^2^) at different timepoints in *APOE4* mice (N=7 mice/group, *p<0.05, **p<0.01, ***p<0.001, ****p<0.0001). E. Two-way ANOVA with multiple comparisons measuring average length of CAA ± SEM per affected vessel in *APOE3* vs. *APOE4* mice at different timepoints (right) (N=7 mice/group, *p<0.05, **p<0.01, ***p<0.001).

### 3D6 acutely increases microglial Aβ_42_ in APOE4 mice

We observed increased levels of Aβ within microglia at early timepoints in *APOE4* mice. A representative image taken from a single z-plane on confocal microscopy demonstrated colocalization of Aβ (immunostained with Moab2, in red) and the intracellular milieu of microglia (marked by the GFP expressed from the CX3CR1 promoter, in green) (Figure 5A). Aβ was seen inside microglia both near and away from a large plaque. To quantify any changes that may have occurred, we performed cell sorting to isolate GFP-fluorescent cells from mouse brains 24-hours following IV injection with 3D6 vs phosphate-buffered saline (Supplemental Figure 3). After lysing those microglia, we measured intracellular Aβ_42_ by ELISA. While levels of intracellular Aβ_42_ did not change in microglia of *APOE3* mice after 3D6 treatment, there was a significant, marked increase of intracellular Aβ_42_ in *APOE4* treated mice compared with control mice (189%, p<0.05) (Figure 5B). These results suggest that Aβ clearance mechanisms observed in *APOE4* mice may be mediated by microglial uptake.

**Figure 5.**
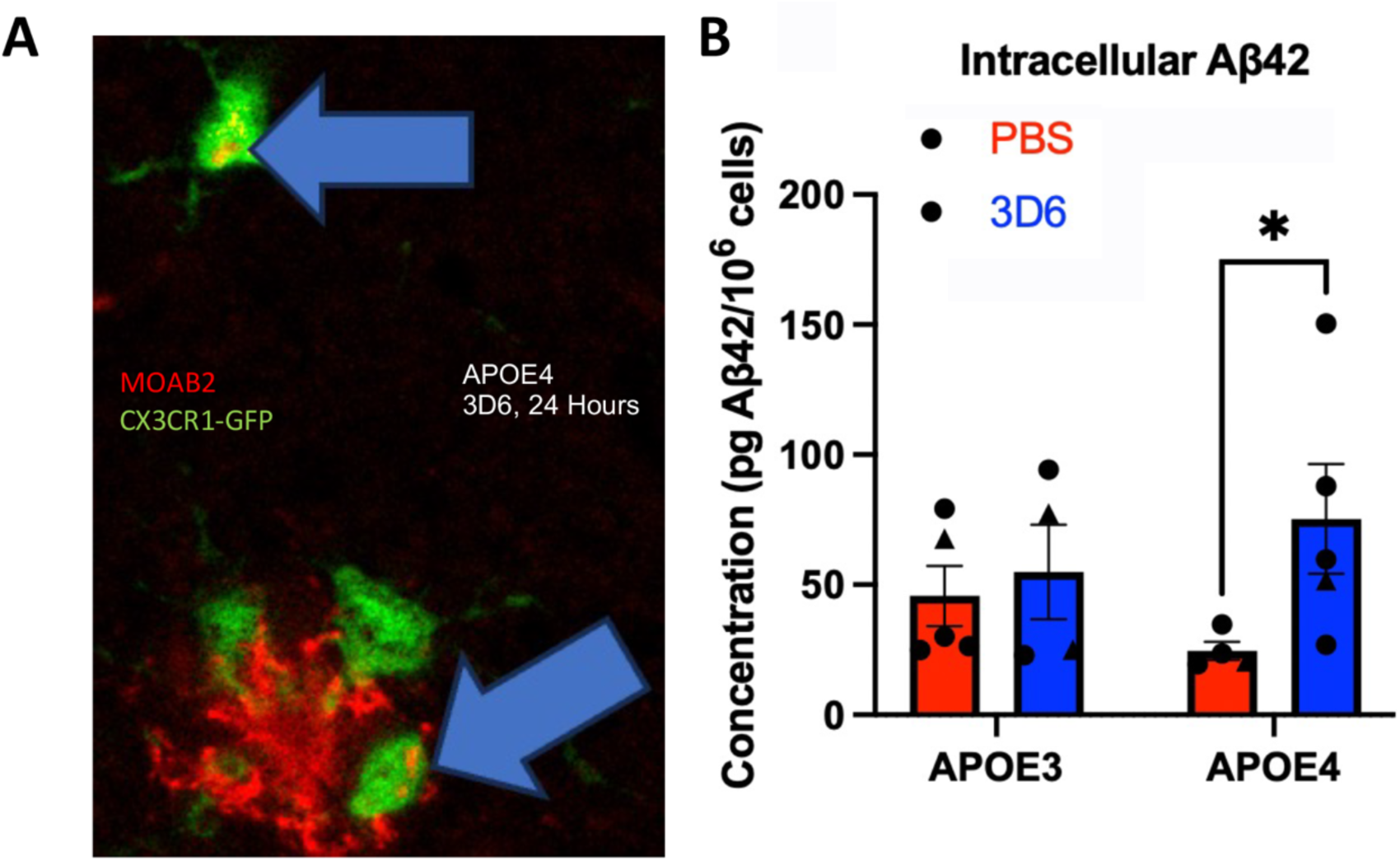
3D6 acutely increases levels of Aβ_42_ located intracellularly in sorted microglia at 24-hours post-treatment preferentially in *APOE4* mice. A. Representative images from 3D6-treated mice at 24-hours post-injection showing Aβ deposits inside microglia both co-localized with a plaque and away from plaques in a single z-plane from confocal image. B. Two-way ANOVA with multiple comparisons measuring mean ± SEM concentration of Aβ_42_ found intracellularly within GFP-sorted cells at 24-hours post-treatment, measured on Aβ_42_ ELISA (N=4-5 mice/group, *p<0.05).

### Acute 3D6 immunotherapy does not increase GFP co-localization with plaques or CAA

We measured co-localization of GFP-fluorescent microglia with both plaques (subiculum) and CAA. Representative images show a 3D6-treated mouse at 10-days post-treatment with a typical microglial response to amyloid (Figure 6A). Under control conditions, about 20% of the deposited Aβ was directly present with the GFP of microglia (Figure 6B). We found no significant changes in GFP co-localization with plaques in either *APOE3* or *APOE4* mice (Figure 6B). Representative images show a roughly 5% co-localization between GFP and a locus of leptomeningeal CAA in a 3D6-treated mouse at day 10 (Figure 6C). In regions with CAA, there was no change in GFP co-localization in *APOE4* mice until day 10, when there was a significant increase in microglial response compared with the earlier treatment timepoints (94-133%, p<0.05) (Figure 6D). These changes in microglia-CAA co-localization were not seen in *APOE3* mice following treatment, showing instead a significant decrease 24 hours post-treatment (47%, P<0.05; Figure 6D).

**Figure 6.**
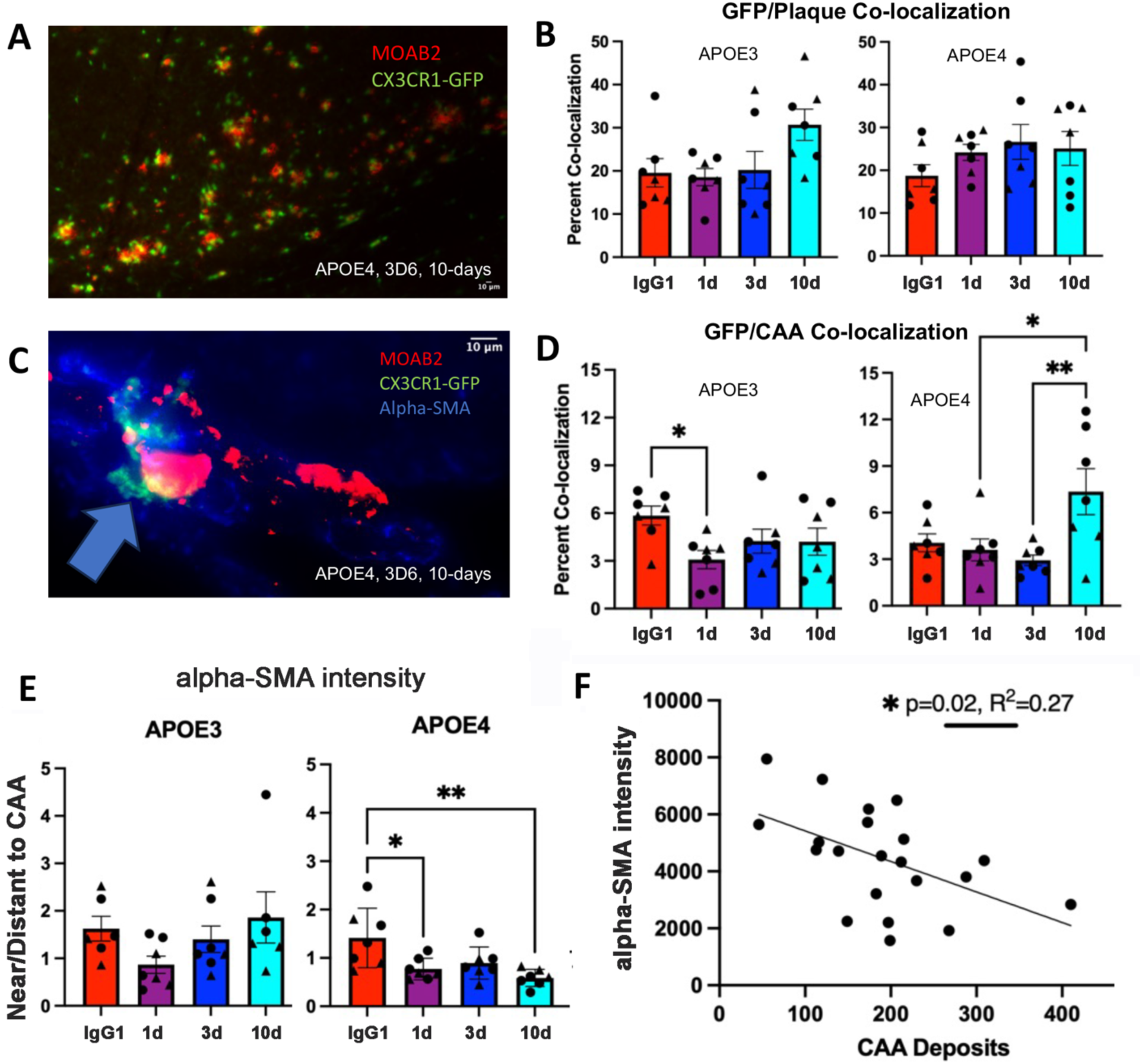
3D6 produces modest changes in GFP co-localization with plaques and CAA with loss of alpha-smooth muscle actin near CAA. A. Representative images showing the microglial response to plaques in a 3D6-treated mouse (10-days). B. One-way ANOVAs with multiple comparisons measuring percent co-localization ± SEM of plaques (Moab2) with GFP in *APOE3* (left) and *APOE4* (right) mice (N=7 mice/group). C. Representative images showing the GFP response to CAA in a 3D6-treated mouse (10-days) (right). D. One-way ANOVA with multiple comparisons measuring percent co-localization ± SEM of CAA (Moab2) with GFP in *APOE3* (left) and *APOE4* (right) mice (N=7 mice/group, *p<0.05, * p<0.01). E. Two-way ANOVA with multiple comparisons measuring mean ratio ± SEM of alpha-smooth muscle actin intensity at regions of CAA compared to regions distant from CAA in *APOE3* and *APOE4* mice using Fiji ImageJ macro (N=7 mice/group, *p<0.05, **p<0.01). F. Simple linear regression measuring correlation of mean number of CAA deposits with average alpha-smooth muscle actin intensity across all *APOE4* 3D6-treated mice (N=7 mice/group, *p<0.05, R^2^=0.27).

### 3D6 decreases alpha-smooth muscle actin near CAA deposits after 10 days

We consistently found that both existing and new CAA in our model forms around alpha-smooth muscle actin (Figure 3A-B). We measured alpha-smooth muscle actin percent area coverage and intensity at the leptomeninges of mice in each condition as an indicator of vessel health. There were no significant changes in overall alpha-smooth muscle actin expression at the leptomeninges after 3D6 treatment in *APOE3* and *APOE4* mice, either in terms of average intensity or area coverage (Supplemental Figure 4A and 4B). We then measured the average intensity of alpha-smooth muscle actin present in deposits of CAA, as well as distant from CAA. We found that 10-days post-treatment, while the baseline ratio is about one (equal alpha-smooth muscle actin away from CAA compared to at CAA), there was a significant 58% decrease in the alpha-smooth muscle actin at the sites of CAA (p<0.01) (Figure 6E). This loss of alpha-smooth muscle actin would indicate that CAA induces degeneration of smooth muscle cells within days of 3D6 treatment. To further test the relationship of CAA with smooth muscle cells, we correlated the number of CAA deposits (Figure 4C) with total average alpha-smooth muscle actin intensity in all *APOE4* treated mice (across timepoints). By linear regression analysis, we found a significant correlation (p=0.02, R^2^=0.27) between increased CAA deposits and decreases in alpha-smooth muscle actin, indicating that higher CAA deposition was associated with loss of smooth muscle cells (Figure 6F). In contrast, for IgG1 control treated-*APOE4* mice, there was no correlation (p=0.92, R^2^=0.002) between CAA deposits and decreased alpha-smooth muscle actin intensity, indicating a 3D6 treatment-specific effect (Supplemental Figure 4C).

### 3D6 increases brain microhemorrhages in APOE4 mice at 10 days

To assess whether our model demonstrates ARIA-H, we performed *in vivo* T1-weighted FLASH MRI and counted the numbers of hypointensities indicative of microhemorrhages throughout the entire brain. We performed MRI scans on five 4-month-old *APOE4* mice at one or two timepoints pre-treatment and at ten days post-treatment with 3D6. We identified distinct points of hypointensity across brain regions under both pre-treatment conditions (yellow arrows) and only post-treatment conditions (blue arrows) conditions (Figure 7A). Ninety-two percent of the lesions observed in pre-treatment scans were also present in post-treatment images. The three mice with two pretreatment measures ten days apart showed no significant change in the number of lesions between pretreatment timepoints. All five mice examined showed similar increases in the number of lesions post-treatment (averaging 32%, p<0.01) (Figure 7B). We tested whether hypointensities demonstrated the presence of bleeds through hemosiderin staining. Brain tissue from three imaged mice was cut into 30 μm sections through regions that demonstrated the hypointensities that we had counted. We stained sections for hemosiderin with Prussian Blue to measure iron deposition indicative of microhemorrhage. We found hemosiderin stains in five of these 30 μm sections and each co-localized with hypointensities identified by MRI (Figure 7C).

**Figure 7.**
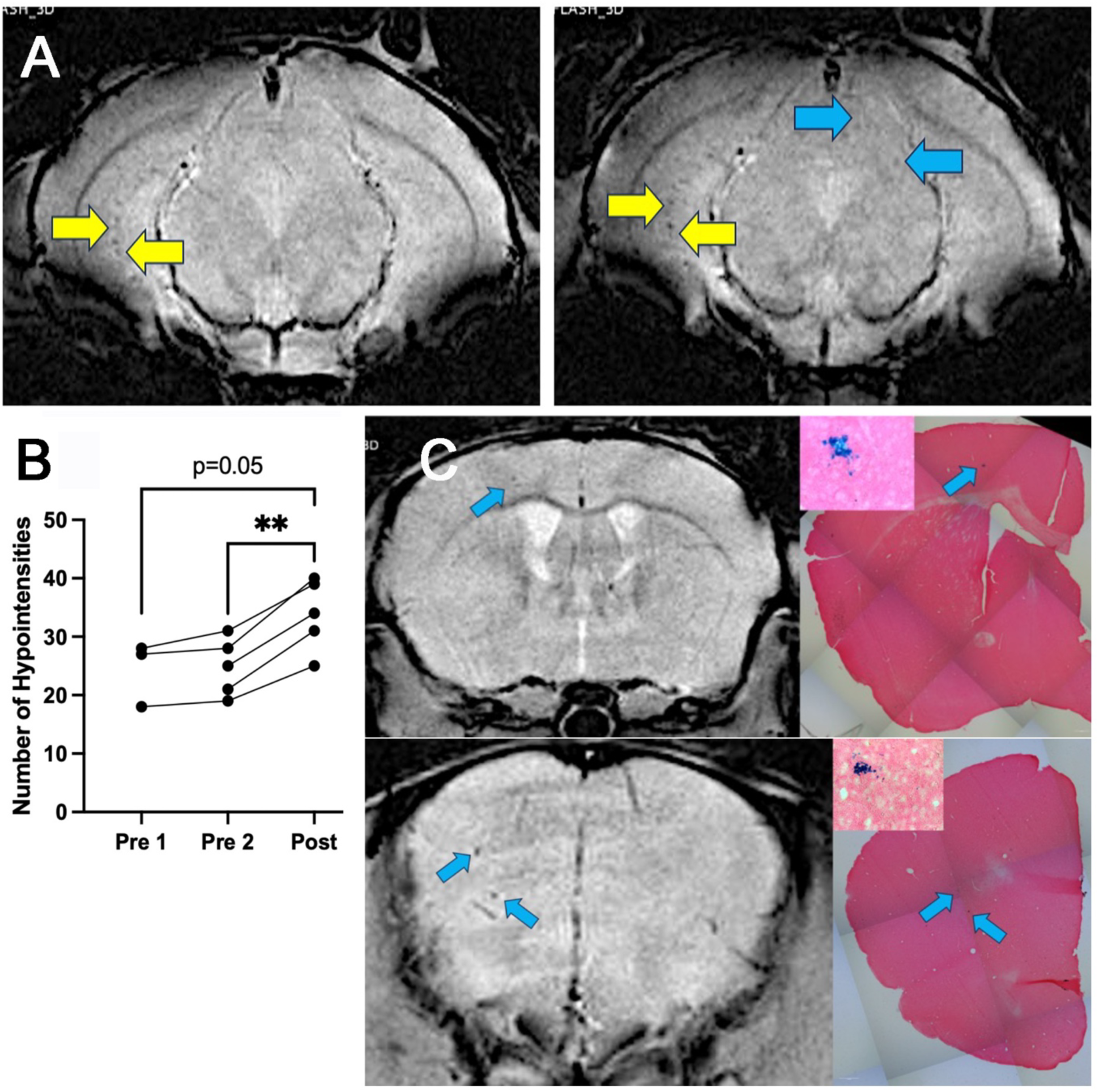
3D6 increases incidence of focal hypointensities 10-days post-treatment in *APOE4* mice. A. Representative image from posterior region of the brain (posterior hippocampus) showing both pre-existing lesions that appear in both images (yellow arrows) and a new lesion that forms post-treatment (blue arrows). B. One-way ANOVA with paired data measuring number of focal hypointensities in each mouse at different timepoints (N=5 mice, *p<0.05, **p<0.01). C. Validation of microhemorrhage as measured on MRI using Perl’s Prussian Blue staining (insets) to detect hemosiderin in analogous brain regions by histology.

## Discussion

Since the first anti-Aβ immunotherapy was FDA approved for the treatment of AD, there have been advances in our understandings of the effects of these drugs ^5^. For example, *APOE4* genotype and dose of antibody have been identified as major risk factors for ARIA ^23^. Potential contributors to ARIA include damages associated with CAA, decreased blood brain barrier integrity, dysregulated neuroinflammation, and higher baseline amyloid load ^5,36^. Our mouse model, incorporating APOE genotype, intravenous exposure to anti-amyloid antibody, and an acute time course of a single exposure addresses novel aspects of CNS damages that contribute to ARIA at times when ARIA incidence is highest ^7,8^. By treating over 80 mice with IV injections of either 3D6 or isotype control, we found that *APOE4* genotype promotes increased CAA, loss of smooth muscle cells, and increased incidence of microhemorrhage, all within one to ten days.

APOE4 genotype is associated with higher levels of amyloid in AD brains ^37,38^. In clinical studies, higher amyloid levels may contribute to elevated rates of ARIA ^36^, and increased rates of plaque clearance may correlate with higher rates of ARIA ^6^. Interestingly, plaque clearance is associated with cognitive improvement in only *APOE4* individuals treated with donanemab ^39^, suggesting a potential *APOE* genotype-mediated role in both the effectiveness and adverse effects. One clinical trial investigating bapineuzumab found differential rates of Aβ (and tau) levels in *APOE4* carriers, with Aβ levels only significantly decreasing in *APOE4* carriers ^40^. We found that there is a rapid clearance of a significant portion of plaques from multiple plaque-rich regions within 10-days of 3D6 immunotherapy in *APOE4* mice. This same degree of clearance does not happen in *APOE3* mice on this timeline. Other preclinical data show clearance of plaques across *APOE* genotypes following chronic antibody treatment ^26,27^. In one cohort of APP/APOE transgenic mice treated chronically with 3D6, there was the largest percent decrease in plaque-load in the *APOE4* group ^41^. The higher level of baseline plaques in *APOE4* mice may contribute to increased Aβ clearance and ARIA risk.

We observed redistribution of amyloid from the parenchyma to the vasculature due to antibody treatment. A clinical study of patients with active anti-Aβ_42_ immunization (in contrast with passive immunotherapy) showed high levels of leptomeningeal Aβ_42_ in a cohort of mostly *APOE4* carriers ^42^. Interestingly, these findings are also associated with increased apoE protein expression in the leptomeninges following treatment, indicative of an APOE-mediated effect ^42^. Severe CAA was also seen in a case study of an *APOE4/4* individual who died of seizures and multiple microhemorrhages after three doses of the anti-amyloid drug lecanemab ^15^. Preclinically, a redistribution of amyloid to the vasculature was seen in the frontal cortex and hippocampus following 2- and 3-months of weekly treatment with an anti-Aβ_28–40_ antibody ^32^. Our data shows that this CAA can form after just one dose of passive immunotherapy in the presence of *APOE4*. This increased CAA correlated with decreased parenchymal Aβ plaques. Given that we also observe increased microhemorrhages on this timeline, we hypothesize that this redistribution of amyloid is one acute mechanism of ARIA.

*APOE4* genotype is linked to higher rates of CAA and cerebrovascular dysfunction ^43^. In mice, vascular amyloid deposits are cleared quickly after exposure to anti-Aβ immunotherapy, but this study used direct application of an anti-amyloid antibody directly to the brain ^44,45^. Rather than decreased CAA after anti-amyloid treatment, we found dramatically higher levels of CAA in *APOE4* mice. Given the established relationship of CAA with spontaneous microhemorrhages in mouse models ^46^, the formation of new loci of CAA following immunotherapy may represent an acute mechanism of ARIA.

Inflammation associated with CAA (CAA-ri) shares clinical characteristics with ARIA ^47^. ARIA related to CAA-ri may be due to deposition of new CAA resulting in vascular damage ^42^, which we found in our studies, or direct interaction of antibody with CAA that triggers a systemic immune response ^48^. The strength of binding of different anti-Aβ immunotherapies to CAA fibrils correlates with rates of ARIA in clinical trials ^49^, indicating that the interaction of antibody with CAA may be partly responsible for moderating risk of ARIA. *APOE4* individuals may have a larger share of antibody-accessible Aβ that could promote ARIA ^50^. It’s important to note that the formation of new CAA and the direct binding of antibody to pre-existing CAA are not mutually exclusive; in fact, the acute formation of new CAA could increase risk of future antibody-CAA interactions with subsequent injections. Future studies with fluorescently conjugated anti-amyloid antibodies (as in Figure 1) would address their direct association with CAA.

CAA formation is linked to the degeneration of smooth muscle cells in both preclinical and clinical models ^51,52^. Chronic treatment with 3D6 in PDAPP mice reduced both the area coverage and intensity of smooth muscle actin ^29^. While the clinical study noted above suggesting redistribution of amyloid from the parenchyma to the vasculature did not exhibit smooth muscle cell loss, it did show splitting of the vessel walls ^42^. We found that smooth muscle actin was damaged at 10-days post-treatment in regions of CAA in *APOE4* mice. We also showed that the increased number of CAA deposits significantly correlated with decreased overall smooth muscle actin intensity. Given the novel deposition of CAA at 24 hours, and then the loss of smooth muscle actin intensity at CAA at 10 days, we conclude that CAA formation quickly causes smooth muscle damage in our *APOE4* mouse model.

There is a rapid increase in microglial cells and processes within three days of passive (IP) Aβ immunotherapy ^53^. When antibody is injected intracranially, there is an immediate microglial response associated with plaque clearance ^54^. Interestingly, beyond three days post-injection, there is continued clearance of plaques without microglial activation in this study, indicating two potential mechanisms of plaque clearance: one microglia-dependent and another microglia-independent ^54^. In these early stages following intracranial injection, microglia may also communicate with other non-parenchymal immune cells to induce a more systemic response ^55^. We observed increased microglial uptake of Aβ at just 24-hours post-treatment, consistent with microglial involvement in the immediate response to immunotherapy. Consistent with the measure of plaque clearance in our independent cohort of mice, intra-microglial Aβ was only seen in *APOE4* mice. *APOE4* microglia have been shown to exist in a more inflammatory state ^56^, perhaps enabling them to more quickly respond to antibody-Aβ complexes. Microglia have also been shown to seed Aβ deposits ^57^; increased seeding could be a cause of elevated levels of plaques in *APOE4* brains. In our work, the increase in intracellular Aβ_42_ at 24-hours corresponding with increased levels of CAA could represent a seeding of CAA by microglia. Microglia have also been shown to clear Aβ via digestive exophagy, an extracellular process mediated by lysosomal enzymes ^58^. The combination of intracellular and extracellular digestion mechanisms of microglia could explain the early uptake of Aβ by microglia and the clearance of plaques, despite limited changes in co-localization of microglia with plaques. Given that there are microglial changes, some of which are *APOE4* specific, that occur following longer-term treatment ^59,60^, we hypothesize that larger-scale immune changes may take a more predominant role later in the treatment course.

The high level of GFP-labeled cells to CAA at 10-days in *APOE4* mice suggests that some immune changes may occur later, perhaps following microhemorrhage. Preclinical studies have established both a microglial response ^61^ and a peripheral immune response ^62^ to microhemorrhage. In human patients, inflammation around CAA contributes to vessel leakage and disruption ^63^. Preclinically, impaired meningeal lymphatic functioning is associated with increased microglial activation following intracisterna magna injection of antibody ^64^. In our work, the poor functioning of these leptomeningeal vessels, perhaps associated with CAA and smooth muscle cell loss, could be responsible for the presence of these GFP-positive cells.

In numerous preclinical models, anti-Aβ immunotherapies increase rates of microhemorrhage as defined by histology ^26,30,65^. MRI is also used to assess ARIA-like markers, and has revealed transient blood brain barrier breakdown and microhemorrhage after chronic treatments ^66,67^. Our study used MRI to measure microhemorrhage following just a single acute IV dose. The relatively stable levels of hypointensities between pretreatment scans, followed by significant increases following treatment, are indicative of a treatment-mediated effect. Given the high baseline levels of hypointensities, we performed hemosiderin staining on three of these mice to test whether there was iron deposition in the same regions as hypointensities on MRI. We consistently found a link between these two measures, providing validation for our MRI model. Together, these data establish that our mechanistic findings of plaque clearance, CAA formation, and smooth muscle actin damage could be linked to increased incidence of ARIA-H after only 10 days. Other imaging approaches would be needed to evaluate the timecourse of ARIA-E induction.

Novel anti-Aβ immunotherapies fused to transferrin receptor antibodies (trontinemab) may improve transport across the blood brain barrier and more safely treat AD patients ^68^. Novel titration schedules of donanemab show that “ramping up” early dosing of immunotherapy can decrease rates of ARIA ^69^. These studies provide evidence that anti-Aβ immunotherapies are not intrinsically tied to ARIA. These developments are consistent with our findings: ARIA risk is highest immediately following treatment initiation and may be linked to vascular damage. We believe that treatment approaches that account for these ARIA risk factors have the potential to treat AD more safely, particularly in the *APOE4* population.

In summary, our model combines *APOE* genotype with the clinically relevant intravenous injections and follows the most immediate effects of this treatment through several approaches. We show that early Aβ clearance, early CAA induction, and the earliest indications of vascular damage occur preferentially in *APOE4* mice. These are previously unrecognized features of amyloid redistribution and ARIA on an acute timeline. The characteristics that we observed in our mouse model will be useful in evaluation of approaches to reduce the different components of ARIA risk, particularly in *APOE4* individuals.

## Methods

### Sex as a Biological Variable

Female and male mice are divided between all treatment groups for all experiments performed. When applicable in quantitative data, female data points are presented as triangles while male data points are presented as circles. Including both female and male mice is of particular importance in AD related studies, as sex differences are observed in rates of AD, although not in incidence of ARIA ^5^.

### Mouse Model

This study was conducted in accordance with ethical standards of the Georgetown University Institutional Animal Care and Use Committee. 5xFAD^+/−^ mice, a mouse model hemizygous for 5 familial AD transgenes (APP K670N/M671L + I716V + V717I, PS1 M146L + L286V), on a C57BL/6J/B6xSJL background, were acquired in house. These mice were crossed with APOE3-KI and APOE4-KI mice to produce E3FAD and E4FAD mice as described previously ^70^. Separately, as previously described ^71^, CX3CR1^GFP/GFP^ mice (JAX stock No. 005582) on a C57BL/6J background (JAX stock No. 000664) were crossed with APOE3-KI (JAX stock No. 029018) or APOE4-KI (JAX stock No. 027894) mice to obtain CX3CR1^GFP/-^ expressing either hAPOE3 or hAPOE4. Finally, these two separate lines were crossed to produce the mouse model: 5xFAD^+/−^, CX3CR1^GFP/-^, APOE3^+/+^ or 5xFAD^+/−^, CX3CR1^GFP/-^, APOE4^+/+^. Mice of the final desired genotypes are shown Figure 1A. 5xFAD mice are heterozygous for viability, while CX3CR1 mice are heterozygous to allow for some expression of CX3CR1. Mice are bred and genotyped in house using PCR assays.

### 3D6 Immunotherapy treatment paradigm

Mice have been divided into 8 groups (N=7-8 each): *APOE3* mice treated with IgG1 isotype control, *APOE3* mice treated with 5mg/kg IV 3D6 (Creative BioLabs cat# PABL-011) euthanized at 24-hours-, 72-hours, and 10-days post-treatment, and *APOE4* mice divided into the same four groups. Four-month-old mice received a single 5mg/kg IV injection of either 3D6 or IgG1 isotype control antibody (Syd Labs cat #PA007126) into the lateral tail vein (Figure 1B). Mice at this age have significant baseline amyloid deposition in characteristic amyloid-rich regions including the subiculum, entorhinal cortex, and thalamus (Figure 1C). Injected solution was in 150μl in phosphate buffered saline (PBS) for all mice. Female and male mice were divided between groups such that each group included at least three mice of each sex. 24 hours-, 72 hours-, or 10-days following injection, mice were euthanized by CO_2_ inhalation and perfused with ice-cold PBS for 1-2 minutes and brains were collected. One hemisphere was fixed in 4% paraformaldehyde (PFA)/4% sucrose solution and transitioned through a sucrose gradient (10%, 20%, and 30% sucrose solutions in PBS) and flash frozen in ice-cold 2-methylbutane. The other hemisphere was snap frozen on dry ice and stored at −80°C for biochemical analyses. The fixed hemisphere was sliced into 30μm sections on microtome for immunofluorescence (IF) experiments (Figure 1B).

### 3D6 Conjugation

To validate that 3D6 reaches the brain, aliquots of 3D6 antibody were conjugated with fluorophore to detect 3D6 in the brain. The ReadyLabel Flex Antibody Labeling Kit (Invitrogen #R10702) was used to conjugate 3D6 to 647nm-fluorophore (3D6-647). The antibody was concentrated and measured in bicinchoninic protein assay to calculate 5mg/kg injection in a 4-month-old *APOE4* mouse of the desired genotype. The mouse was euthanized at 72-hours post-injection as described above, and fluorescent and confocal imaging showed 3D6-647 reaches the brain and reacts with plaques (Figure 1E).

### Immunofluorescence

One brain hemisphere was fixed in 4%PFA/4% sucrose solution, dehydrated, and flash frozen as described above, and stored in cryoprotectant. Sections (4 hemispheres per mouse per assay) were washed in PBS (pH 7.4) with antigen retrieval in cold citrate buffer (pH 6.0) before being blocked in 10% normal goat serum (NGS) and 5% bovine serum albumin (BSA) for 2 hours with gentle shaking. Brain tissue was treated overnight at 4°C with gentle shaking in primary antibody: anti-mouse Moab-2 antibody (NovusBio cat# NBP2-13075) to stain Aβ @1:500 dilution and anti-rabbit alpha smooth muscle actin antibody (Abcam cat# ab124964) @1:250 dilution, each in PBS with 1% NGS, 1% BSA, and 0.1% tween-20. After being washed 3 times in PBS, tissue was then treated in secondary antibody: anti-mouse IgG2b AlexaFluor350 (Invitrogen cat# A-21140) @1:1000 dilution and anti-rabbit AlexaFluor594 secondary antibody (Invitrogen cat# A-11012) @1:2000 dilution in 1% NGS for 2 hours at room temperature with gentle shaking. Secondary antibody against mouse IgG2b (rather than general anti-mouse secondary) was used to stain for Moab2 to limit staining of endogenous mouse antibody.

Images were taken at consistent exposure on Zeiss microscope: Moab2 (Aβ) was measured in DAPI channel (350nm), GFP (microglia) was measured in eGFP channel (488nm), and alpha-smooth muscle actin (cerebral arteries/arterioles) were measured in AlexaFluor594 channel (594nm). 4 brain slices containing hippocampus (subiculum), entorhinal cortex, and thalamus were analyzed for each mouse for each experiment. Images were taken at 10x (for plaque counting and area coverage analysis, microglia analysis) and 20x magnification (for CAA and alpha-smooth muscle actin analysis). Images were analyzed on Fiji ImageJ software for total number of plaques and percent area coverage of Aβ. To measure size of plaques, we used the Analyze Particles tool on Fiji ImageJ and divided bins into 5-25, 26-100, 101-200, 201-600, 601-1000, and 1000+ μm^2^. Deposits in the 5-25 μm^2^ group were counted in the plaque size analysis, but not for plaque number. This tool incorporates “circularity” of deposits such that only plaques are counted; we adjusted circularity to 0.05-1.00. Plaque number counts were validated by hand counting. Regions where Moab2 surrounded alpha-smooth muscle actin were counted as loci of CAA, such that each loci represents an individual vessel impacted by CAA. The number of CAA deposits were counted differently because multiple CAA deposits could form on a single vessel. Regions of interest were drawn around these vessels to measure the length of CAA within a vessel. As with plaques, to measure size of individual CAA deposits the Analyze Particles tool was used, this time with size bins of 5-10, 11-25, 26-50, 51-100, 101-200, and 200+ μm^2^. Areas of co-localization between GFP and Moab2 were measured using the JACOP plug-in on Fiji ImageJ. Plaque co-localization was measured from the subiculum, while CAA co-localization was measured from the leptomeninges. At least 4 images were analyzed per each section, per mouse (at least 16 images analyzed per mouse per analysis). All experimenters were blinded to the group and genotype during all analysis.

### Confocal Imaging

Confocal Z-stacks for certain representative images were taken with a laser scanning microscope system (Thor Imaging System Division) equipped with 488/561/642 nm laser and Green/Red/Far-red filters and mounted on an upright Elipse FN1 microscope (Nikon Instruments). Images were assessed both in single z-planes to confirm intramicroglial Aβ and maximum intensity projection for more general imaging.

### Cell Sorting

For cell sorting, hemi brains were sectioned into six sagittal slices using a surgical scalpel in a petri dish immediately following excision. Cells were then dissociated from the neural tissue using the MACS Adult Brain Dissociation Kit according to manufacturer’s instructions (Cat #130-107-677, Miltenyi Biotec). Cells were dissociated from the tissue in 2mL within a gentleMACS C-tube in a gentleMACS Octo-Dissociator at 37 °C for 30 minutes. Following enzymatic dissociation, the single-cell suspension was processed for debris removal using the debris removal solution from the MACS Adult Brain Dissociation Kit. Samples were centrifuged and decanted, and the cell pellet was suspended in 500 μl of cold Dulbecco’s-PBS. Samples were kept on ice until sorting. Cell sorting was conducted by the Georgetown University Flow Cytometry and Cell Sorting Shared Resource. For sorting, sytox blue viability dye was used to differentiate dead cells from live ones. GFP-positive cells were isolated from the viable cell population and sorted for use. Total cell number per sorted population was recorded, and representative data was captured for the first 100,000 events. Following sorting, GFP-positive cells were stored in 1mL PBS in polypropylene FACS tubes, and frozen at −20°C until cell lysis/protein analysis.

### ELISA

For sorted GFP-positive cells, ELISA was performed using Human Amyloid beta (aa1-42) Quantikine ELISA Kit to measure Aβ42 (Biotechne R&D Systems cat #DAB142). Cell sorted samples were prepared with RIPA buffer and protease-phosphatase inhibitors. Samples were then diluted 1:1 in diluent buffer provided in kit before being run on ELISA. Samples were normalized to the total number of cells collected via FACS cell sorting.

### Measurement of alpha-smooth muscle actin concentration near or distant to CAA

A macro was developed on Fiji ImageJ to measure the average intensity of alpha-smooth muscle actin at loci of CAA and distant from CAA. Beginning with z-stack images with separate channels for Moab2 (Aβ) and alpha-smooth muscle actin, the macro produced average intensity projections. We drew a region of interest around the leptomeningeal region containing CAA. The macro created masks within the region of interest that defined regions of CAA and regions of alpha-smooth muscle actin. The macro produced values for the alpha-smooth muscle actin intensity in total, and the alpha-smooth muscle actin intensity within individual masks of CAA. The average alpha-smooth muscle actin intensity at CAA was subtracted from the overall average alpha-smooth muscle actin intensity to produce an approximation for the intensity away from CAA. A ratio was calculated that measured the average intensity of alpha-smooth muscle actin at regions of CAA compared to distant from CAA.

### T1-weighted FLASH MRI

Longitudinal *in vivo* MRI was performed using the Bruker 7-tesla/30 AVANCE NEO MR imager run by ParaVision 360 software, using a high resolution 12 cm gradient in the Georgetown Preclinical Imaging Research Laboratory (PIRL, S10 OD034435). Anesthetized (1.5% isoflurane in oxygen) mice were placed in a custom-manufactured (ASI Instruments, Warren, MI) stereotaxic device with built-in temperature and cardio-respiratory monitoring, as previously described ^72,73^ compatible with a 23 mm transmit-receive mouse brain coil. To detect cerebral microbleeds, a T1-weighted FLASH (Fast Low Angle Shot) sequence was run with TR: 500 ms, TE: 8.0 ms, FA: 40, FOV: 18 x 18 mm, Matrix: 256 x 256 and slice thickness/distance: 0.6 x 0.6 mm. This technique enables the visualization of hemorrhagic hypointense foci which represent focal deposits of iron or hemosiderin in the brain. Mice were imaged either once or twice before treatment (each scan 10 days apart) and once 10-days post-treatment. Lesions were counted as parenchymal microhemorrhages if they were non-symmetrical, non-continuous between slices, not directly adjacent to the surface of the brain, and away from any clear MRI artifacts. Raters were blinded to information about which timepoint represented each MRI scan. Differences in the numbers of lesions between timepoints were measured using one-way ANOVA with pair-matching, with multiple comparisons.

### Perl’s Prussian Blue Staining

This protocol was adapted from the Iron Stain Assay Procedure (Abcam 150674). After MRI imaging for microhemorrhage, tissue was sectioned coronally at 30 μm. Free floating tissue sections rinsed in 1mL of deionized water (dH2O) for three 5-minute washes. Tissues sections were then stained for iron with Prussian Blue Stain. The sections were incubated with equal amounts of Potassium ferrocyanide solution and Hydrochloric Acid Solution for 20 minutes. The tissues were washed with dH2O for 5 minutes followed by a 5-minute counterstain with a 1:20 dilution of the Nuclear Fast Red Solution (5%). Tissues were then washed four times in 10-minute increments with dH2O. Dehydration of tissue was performed with two washes of alcohol, firstly with 95% alcohol followed by absolute alcohol. Tissue was cleared and mounted on slides with a DPX mounting media. The hemosiderin staining was viewed under a microscope to confirm that hemosiderin localized at the same areas of the previously rendered MRI bleeds.

### Statistics Analyses

Data in all graphs are displayed as mean ± SEM. For each graph each data point represents a mouse, and when they are differentiated, each triangle represents a female mouse, and each circle represents a male mouse. Data are analyzed using student’s t-test, one-way ANOVA, or two-way ANOVA, with corrections. All tests are two-tailed. For statistical significance: * p<0.05, ** p<0.01, *** p<0.001, **** p<0.0001. Sidak’s multiple comparisons were used. Outlier tests were performed using ROUT. Experimenters were blinded to treatment group and timepoint for all analysis. Statistical analyses were all done using GraphPad Prism 10.0.

### Study Approval

All studies were carried out following the Guide for the Care and Use of Laboratory Animals as adopted by the U.S. National Institute of Health and approved by Georgetown University Animal Care and Use Committee, approval protocol 2016–1160.

### Data Availability

All primary datasets and original figures can be provided upon request.

## Author Contributions

PP wrote the manuscript, designed research studies, conducted experiments, and acquired and analyzed data. GH conducted experiments and acquired data. EX analyzed data. GS conducted experiments and provided analytical techniques. NS analyzed data. YL conducted experiments. CA conducted experiments. OR conducted experiments. RST edited manuscript and provided feedback. GWR designed research studies, conceived of manuscript scope, and served as principal investigator of the studies. All authors provided edits for the manuscript.

## Funding Support

NIH T32AG071745, Aging and Alzheimer’s Research Training Program.

NIH S10OD034435 (Albanese)

P30 CA051008 (Weiner)

## Acknowledgements

The authors thank all members of the Rebeck Lab past and present for support and assistance. We especially thank former lab manager Christi Anne Ng for training in animal handling skills and IV injections. We thank the Preclinical Imaging Research Laboratory and the Georgetown Division of Comparative Medicine (DCM) husbandry and vet technician staff for assistance with animal work. We thank the Georgetown University custodial staff and environment health and safety staff for ensuring our labs remain clean and safe.

---

**Supplemental Figure 1.**
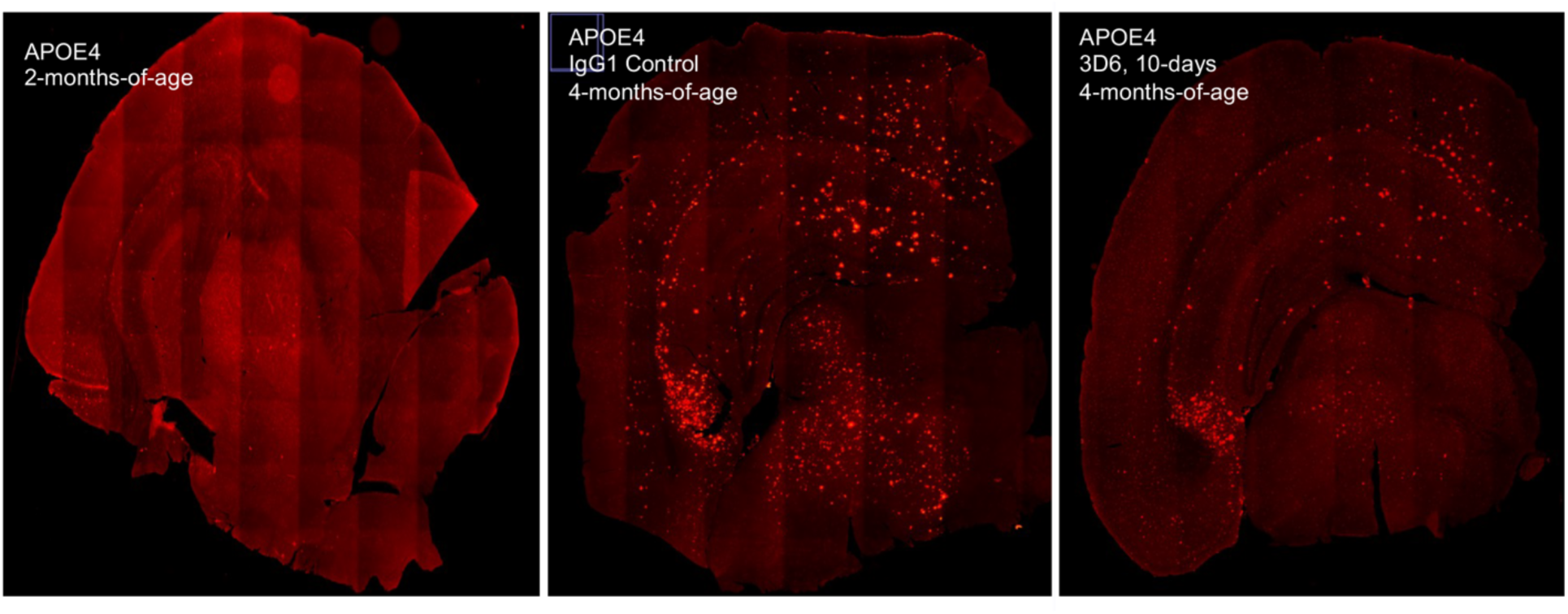
Aβ Plaques are widespread in our mouse model by 4 months of age and a portion of those plaques are cleared 10-days following 3D6 treatment. Plaques were stained in a 2-month-old male 5xFAD^+/−^, CX3CR1^GFP/-^, APOE4^+/+^ mouse (left) and 4-month-old male 5xFAD^+/−^, CX3CR1^GFP/-^, APOE4^+/+^ mouse (middle) using Moab-2. Very early plaque accumulation was seen at 2 months, with widespread plaques developing by 4-months-of-age. Ten days following 3D6 treatment, a decreased distribution of plaques was seen in a male 5xFAD^+/−^, CX3CR1^GFP/-^, APOE4^+/+^ also at 4-months-of-age (right).

**Supplemental Figure 2.**
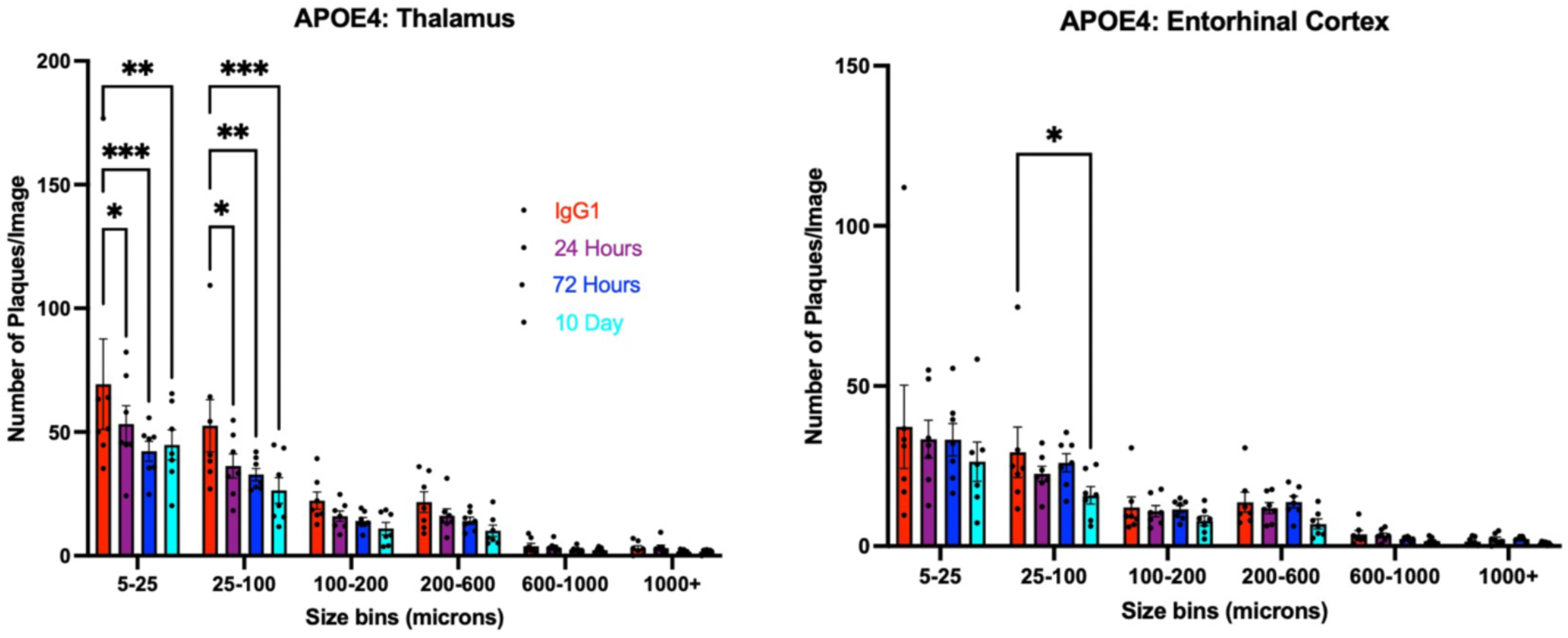
Plaques of smaller sizes are cleared in the thalamus, with limited clearance in any size bin in the entorhinal cortex. Two-way ANOVA with multiple comparisons measuring mean ± SEM number of plaques/image in different size bins of Aβ plaques in the thalamus (left) and entorhinal cortex (right) of *APOE4* mice. Bins are 5-25 μm^2^, 25-100 μm^2^, 100-200 μm^2^, 200-600 μm^2^, 600-1000 μm^2^, and over 1000 μm^2^ (N=7 mice/group, *p<0.05, **p<0.01, ***p<0.001, ****p<0.0001).

**Supplemental Figure 3.**
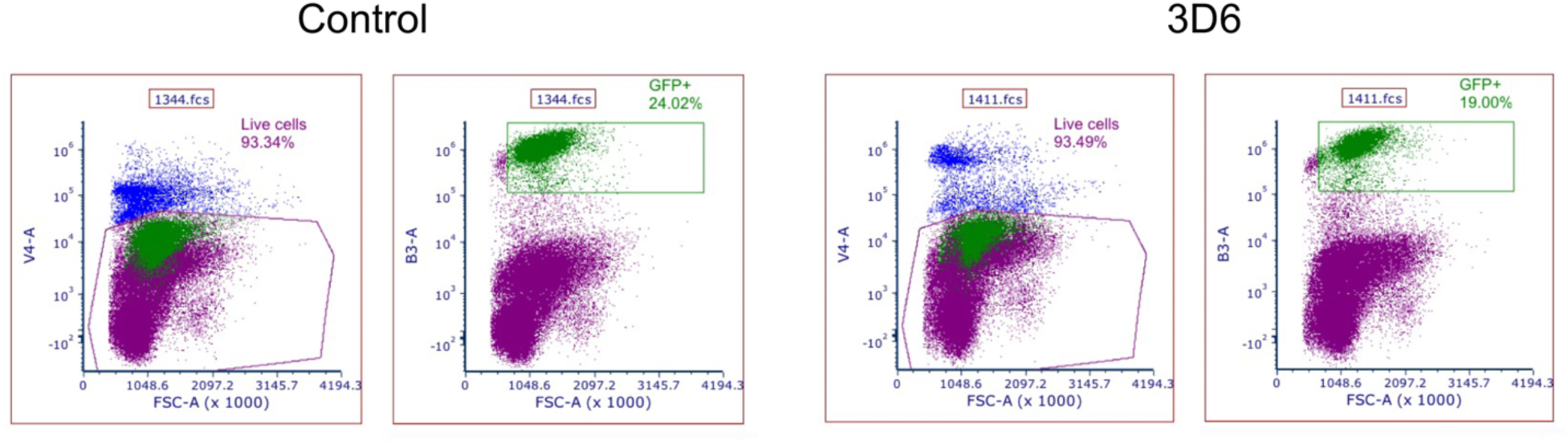
Representative plots showing cell sorting process to isolate GFP-positive cells. GFP-positive cells were isolated first by isolating the live cell population (blue dots) and then the GFP-positive population (green dots). A representative example for a control (PBS) mouse and 3D6-treated mouse at 24-hours post-treatment are shown.

**Supplemental Figure 4.**
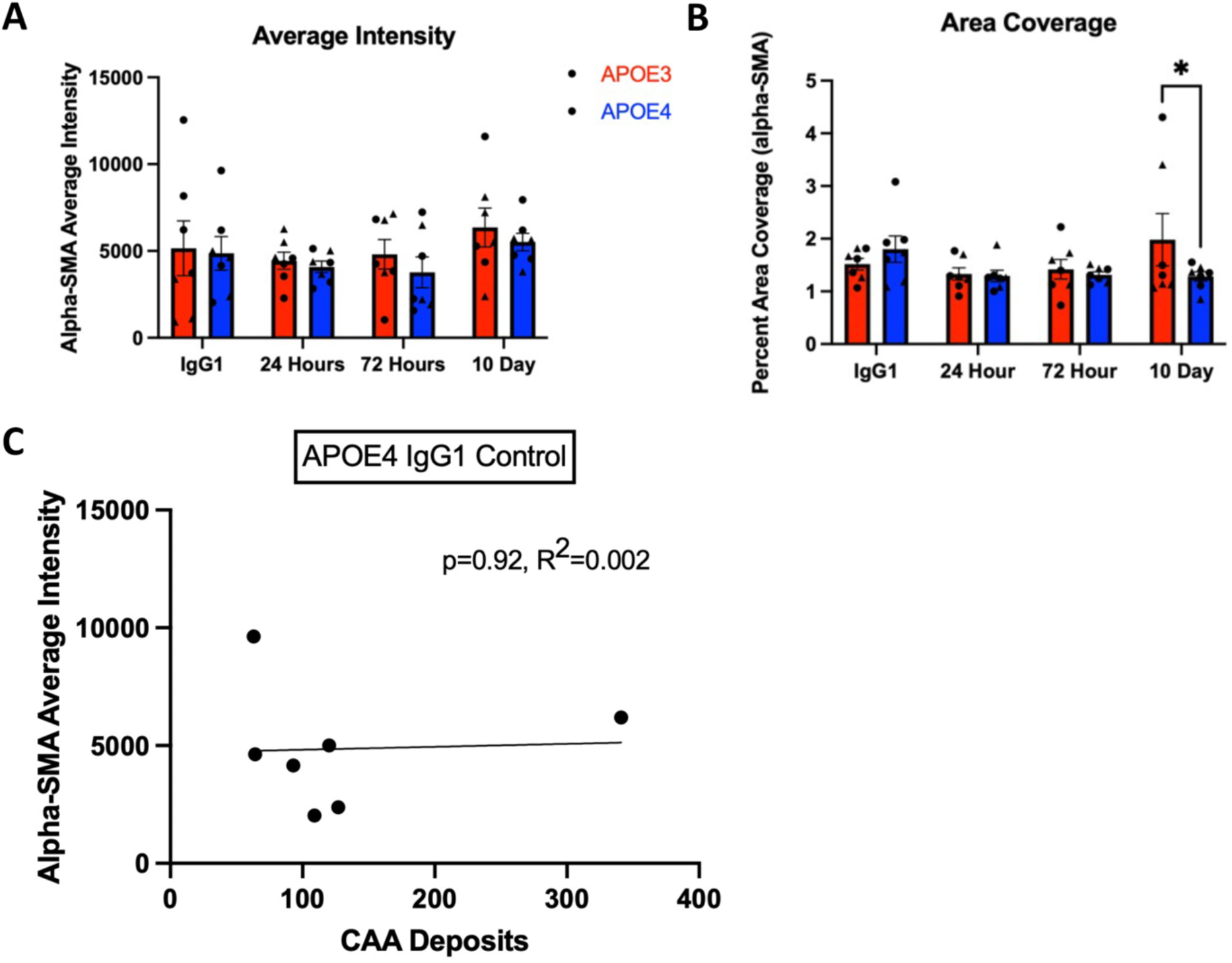
Increases in CAA deposits do not correlate with any change in alpha-smooth muscle actin intensity in IgG1 control mice. A-B. Two-way ANOVA with multiple comparisons measuring mean ± SEM number of alpha-smooth muscle actin intensity (A) and percent area coverage (B) in the leptomeninges of *APOE3* vs. *APOE4* mice at different points (middle) (*p<0.05). C. Simple linear regression measuring correlation of mean number of CAA deposits with average alpha-smooth muscle actin intensity across *APOE4* IgG1 control-treated mice (N=7 mice/group, *p=0.92, R^2^=0.002).

